# PathDSP: Explainable Drug Sensitivity Prediction through Cancer Pathway Enrichment

**DOI:** 10.1101/2020.11.09.374132

**Authors:** Yi-Ching Tang, Assaf Gottlieb

**Affiliations:** Center for precision health, School of Biomedical informatics, University of Texas Health, Science Center at Houston, Houston, TX, 77030

**Keywords:** Anticancer drug response prediction, Drug sensitivity prediction, Precision cancer medicines

## Abstract

Computational approaches to predict drug sensitivity can promote precision anticancer therapeutics. Generalizable and explainable models are of critical importance for translation to guide personalized treatment and are often overlooked in favor of prediction performance.

Here, we propose a pathway-based model for drug sensitivity prediction that integrates chemical structure information with enrichment of cancer signaling pathways across drug-associated genes, gene expression, mutation and copy number variation data to predict drug response on the Genomics of Drug Sensitivity in Cancer (GDSC) dataset. Using a deep neural network, we outperforming state-of-the-art deep learning models, while demonstrating good generalizability a separate dataset of the Cancer Cell Line Encyclopedia (CCLE) as well as provide explainable results, demonstrated through case studies that are in line with current knowledge. Additionally, our pathway-based model achieved a good performance when predicting unseen drugs and cells, with potential utility for drug development and for guiding individualized medicine.

## Introduction

Tailoring drugs to patients based on their genomic and environmental factors is one of the ultimate goals of precision medicine. Within precision oncology, using the molecular signatures of targeted genes has been useful for targeted therapy (Baudino, 2015). The use of machine learning algorithms have significantly progressed in predicting drug response by integrating genetic features and chemical structure information. Menden et al. derived multiple features from cell lines, including microsatellite instability status, mutations and copy number variations to predict drug response, and demonstrating that that chemical information improved significantly the drug models (Menden *et al,* 2013). Another work, by Wang et al., used matrix factorization to predict drug response from low dimensional drug similarity space and cell line similarity space (Wang *et al,* 2017) while a work by Liu et al. performed better by including drug response similarity (Liu *et al,* 2018). Recently, Li et al. (Li *et al,* 2019) published a method called DeepDSC that predicts drug response by encoding gene expression features through an autoencoder and feeding the encodings together with chemical structure information into a deep neural network.

While these computational models attempt to improve performance, the focus on explainable and generalizable models remains limited. Being able to explain the model results to oncology researchers can diminish a significant barrier for translation of prediction models to clinical setting of precision oncology. In order to address this barrier, a study by Yang et al. (Yang *et al*, 2018) built a Bayesian model for inferring the relation between drug target proteins and cancer signaling pathway activities. This work assumed that if a drug pathway is activated in a tumor, then the tumor cells are likely to be sensitive to drugs that target genes in that pathway. Mathematically, the drug response variable (e.g., IC50) is factorized into drug target feature and signaling pathway feature. While gaining interpretability, this approach displayed low performance.

In this study, we integrated ideas of cancer pathways with deep learning constructs to develop a pathway-based model for the prediction of drug sensitivity in cancer, called PathDSP, which both performs well and is also explainable. The rationale is that drugs exert their therapeutic effects by affecting target proteins, further signaling downstream pathways. Activation of signaling pathways thus may indicate whether cells are sensitive or resistant to a drug when the activated signaling pathways are important for cell growth or death. We integrated drug-based pathway enrichment scores across 196 cancer pathways (Schaefer *et al,* 2009) with cell-based pathway enrichment scores in these pathways. Testing our model on the Genomics of Drug Sensitivity in Cancer (GDSC) (Yang *et al,* 2012) and the Cancer Cell Line Encyclopedia (CCLE) databases (Barretina *et al*, 2012), we demonstrate better performance than previously published deep learning approaches while case studies show that pathway features agree with current knowledge on drugs’ mechanism of action. To the best of our knowledge, this is the first pathway-based deep neural network for drug sensitivity prediction and it provides a flexible framework to incorporate additional pathways using prior knowledge. Demonstrating the generalizability of our approach across databases and for predicting response to new drugs and cell lines, our approach can thus be useful both for drug development and for guiding individualized medicine.

## Results

### Model selection

We designed PathDSP for predicting drug response in cancer cell lines based on the rationale that cancer pathways would represent well drug therapeutic effects (Figure 1). We predicted drug responses for 153 drugs across 319 cell lines assembled from the Genomics of Drug Sensitivity in Cancer (GDSC) with available gene expression, somatic mutation and copy number variations data (Methods). Our prediction scheme included predicting of the response values (log-transformed IC50) based on two drug-based data types and three cell line-based data types. The drug-based features include chemical structure molecular fingerprints (from here on referred as CHEM) and pathway enrichment of gene subnetwork of drug-associated genes (Drug-Gene Network, DG-Net). The cell-line-based features include pathway enrichment scores for gene expression (EXP), mutation (MUT-Net) and copy-number variation (CNV-Net) data (Methods).

**Figure 1:**
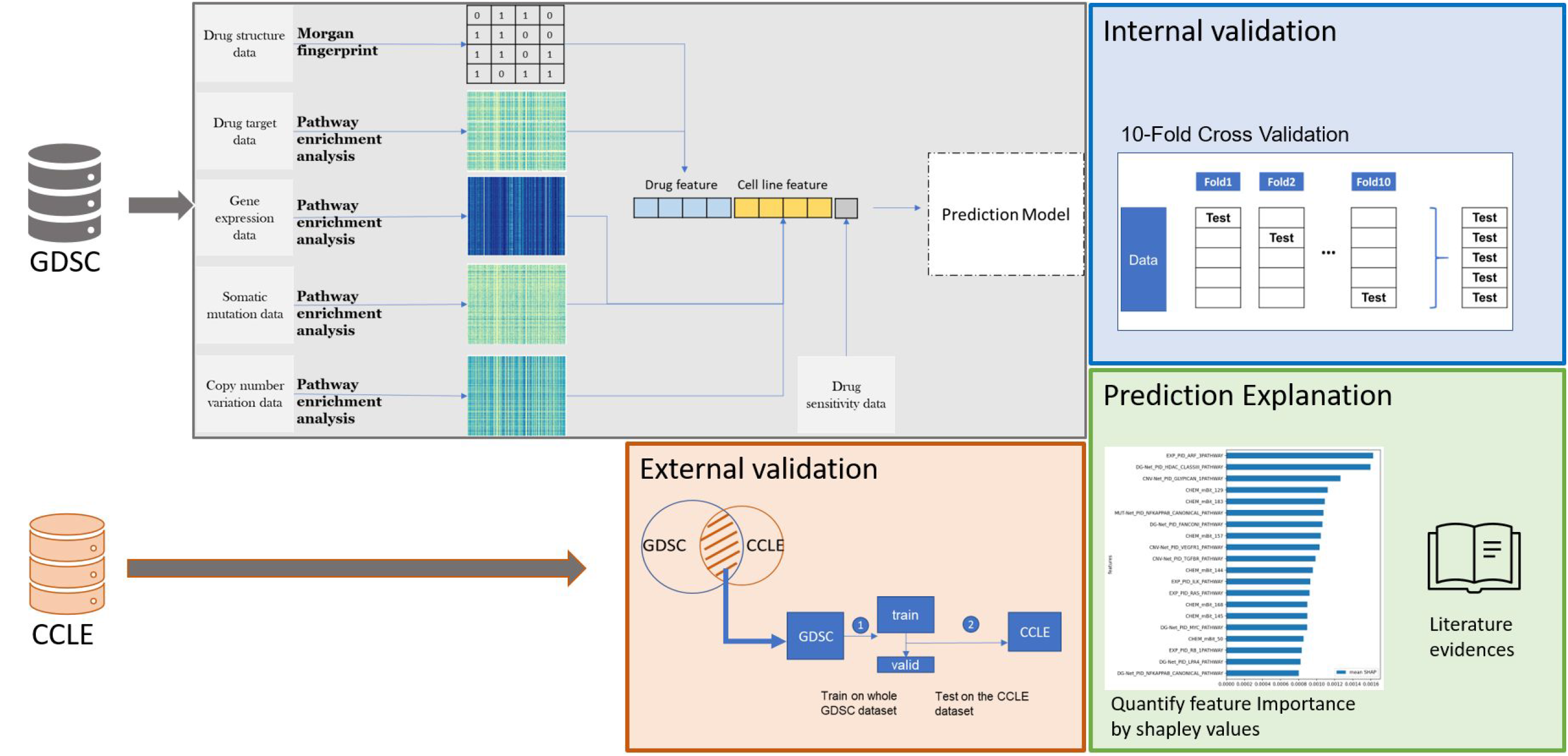
Illustration of the PathDSP method and validation plan.

We compared the performance of six machine learning algorithms, including ElasticNet, CatBoost (Prokhorenkova *et al,* 2018), XGBoost (Chen & Guestrin, 2016), Random Forest (Breiman, 2001), Support Vector Machine (SVM) (Chang & Lin, 2011), and the fully connected neural network (FNN). The result of 10-fold cross validation demonstrated that FNN obtained the best performance with a RMSE of 0.35±0.02 (Table S1). Therefore, we used the FNN model throughout the study.

For the FNN model, we further compared performance of subsets of the data types. Unsurprisingly, the combination of all data types obtained the lowest RMSE of 0.35±0.02 (Table 1). Using either CHEM, EXP, MUT-Net or CNV-Net as single drug or cell-related data types performed very similar (0.42<RMSE<0.44), while DG-Net without CHEM performed worst (RMSE=0.54). We thus used this best performing model for the following analysis.

**Table 1.** The RMSE of 10-fold cross validation obtained by including different subsets of the data types.

| Prediction Scheme | Drug Feature | Cell Feature | RMSE | R<sup>2</sup> | PCC |
| --- | --- | --- | --- | --- | --- |
| Drug-oriented | CHEM | EXP + MUT-Net + CNV-Net | 0.42 | 0.87 | 0.93 |
|  | DG-Net | EXP + MUT-Net + CNV-Net | 0.54 | 0.8 | 0.9 |
| Cell-oriented | CHEM + DG-Net | EXP | 0.43 | 0.87 | 0.93 |
|  | CHEM + DG-Net | MUT-Net | 0.42 | 0.88 | 0.94 |
|  | CHEM + DG-Net | CNV-Net | 0.44 | 0.86 | 0.93 |
| Reference Line | CHEM + DG-Net | EXP + MUT-Net + CNV-Net | 0.35 | 0.92 | 0.96 |

### Comparison with other methods

We compared PathDSP with four previous studies, including DNN by Menden et al, SRMF, NCFGER, and DeepDSC (Menden *et al,* 2013; Wang *et al,* 2017; Liu *et al,* 2018; Li *et al,* 2019). All models used the same drug response and provided RMSE on the GDSC dataset. PathDSP outperformed these four models, where the next best performer was DeepDSC with an RMSE of 0.52, followed by other models with RMSE between 0.83 and 1.43 (Figure 2).

**Figure 2:**
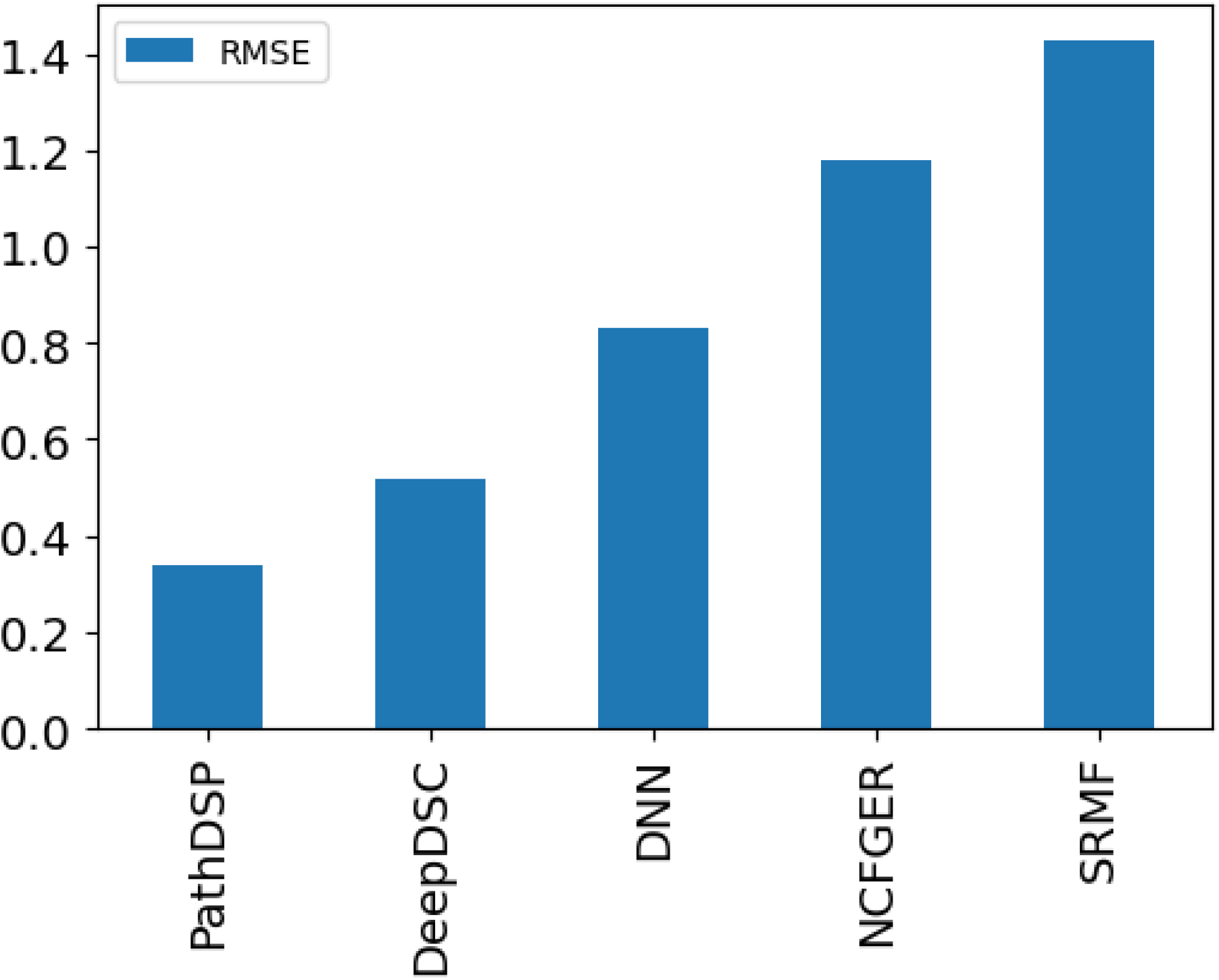
Comparison of PathDSP performance relative to other methods.

### Prediction of new drugs and cell-lines

*In-silico* inference of drug response to a new experimental molecule would be beneficial for pharmaceutical research and drug design, reducing the high cost of large-scale drug screening tests. Predicting drug response for new cell lines, on the other hand, could be translated to clinical setting when oncologists are tasked with prioritizing treatment options for a new patient, potentially translating the cell-line genomic data to the patient’s genomic data. To address these scenarios, we performed leave-one-drug-out (LODO) and leave-one-cell-out (LOCO) by removing one drug or cell line, respectively, from training and assessing the prediction error for that missing drug or cell line (Methods). We obtained an average RMSE of 0.98±0.62 from LODO, and an average RMSE of 0.59±0.17 from LOCO. In particular, our LODO model obtained lower RMSE than DeepDSC, the only model that performed leave-one-drug-out out of the four previously compared models (1.24±0.74).

### Generalizability of PathDSP

We further tested the generalizability of PathDSP across different datasets. We applied the model trained on the GDSC dataset to an independent dataset from the Cancer Cell Line Encyclopedia (CCLE) (Barretina *et al,* 2012). PathDSP obtained an RMSE of 1.15 when tested on the entire dataset of CCLE and when further validated only on the set of drug-cell line pairs shared between GDSC and CCLE, our model obtained an RMSE of 0.95. Notably, the difference in the drug response measurements between the two datasets makes this task challenging. While GDSC provide a large range of IC50 values, CCLE used incomplete response curves, capped at a maximal tested concentration of 8 μM (Haibe-Kains *et al,* 2013; Pozdeyev *et al,* 2016), resulting in different response values of the same drug-cell line combinations across the two datasets (Figure 3). To demonstrate the difference between the datasets, we computed the expected RMSE if PathDSP had perfectly modeled the training data (GDSC). Computing the RMSE of the true response values of GDSC and testing against the response values of CCLE on the shared subset of drug-cell line pairs, the result is higher than the one we obtained with our algorithm (1.13 RMSE vs. our predicted 0.95 RMSE), suggesting that PathDSP is able to generalize well. Notably, despite different distributions of input features between the two datasets (e.g. expression measurements), the pathway-enriched features of PathDSP displayed more consistent patterns between two datasets (Figure 4). These results suggest with more consistent IC50 measurements, PathDSP has the potential to obtain better performance.

**Figure 3:**
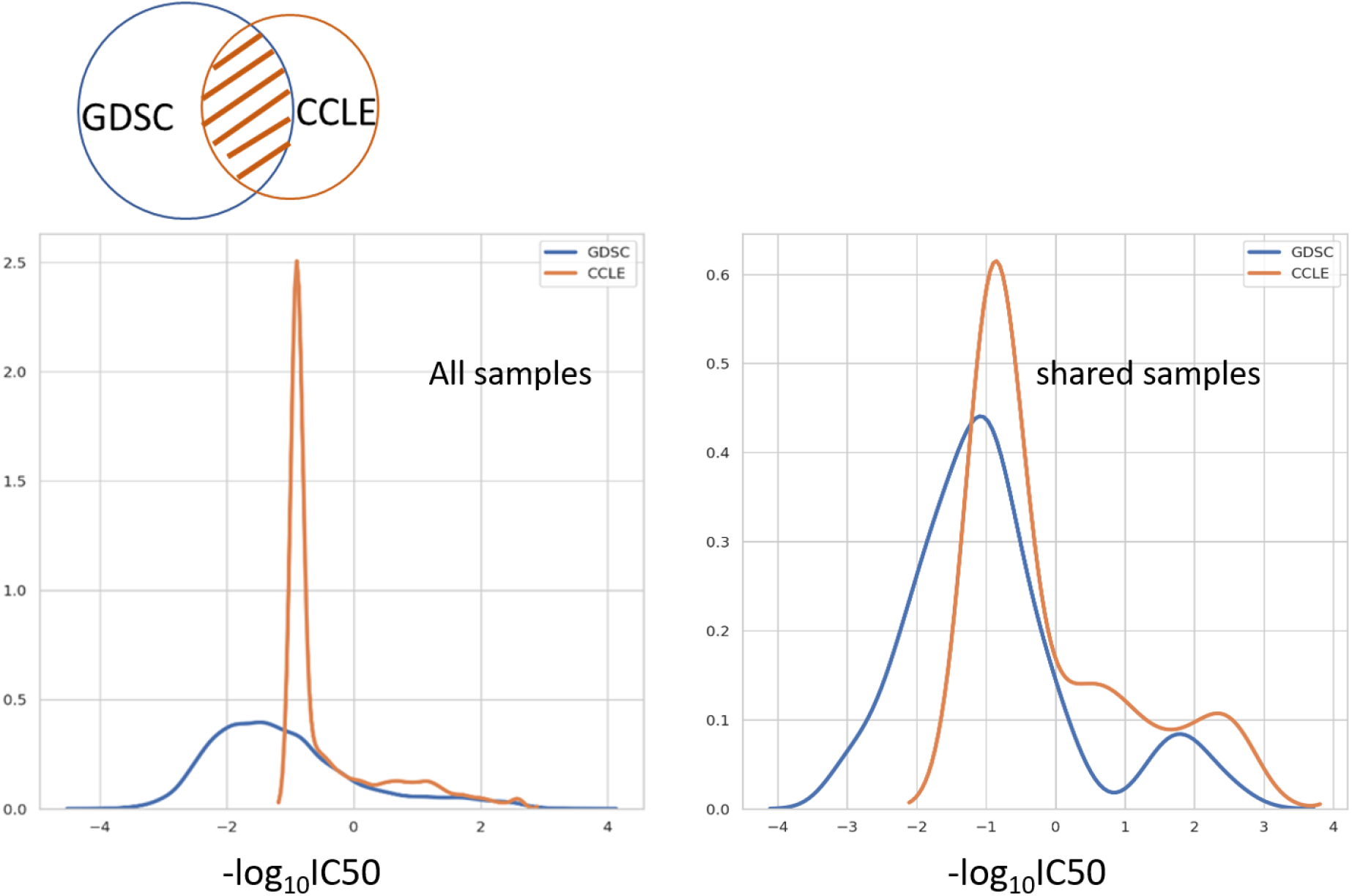
Comparison of drug response distribution between the GDSC (blue line) and CCLE (orange line) databases.

**Figure 4:**
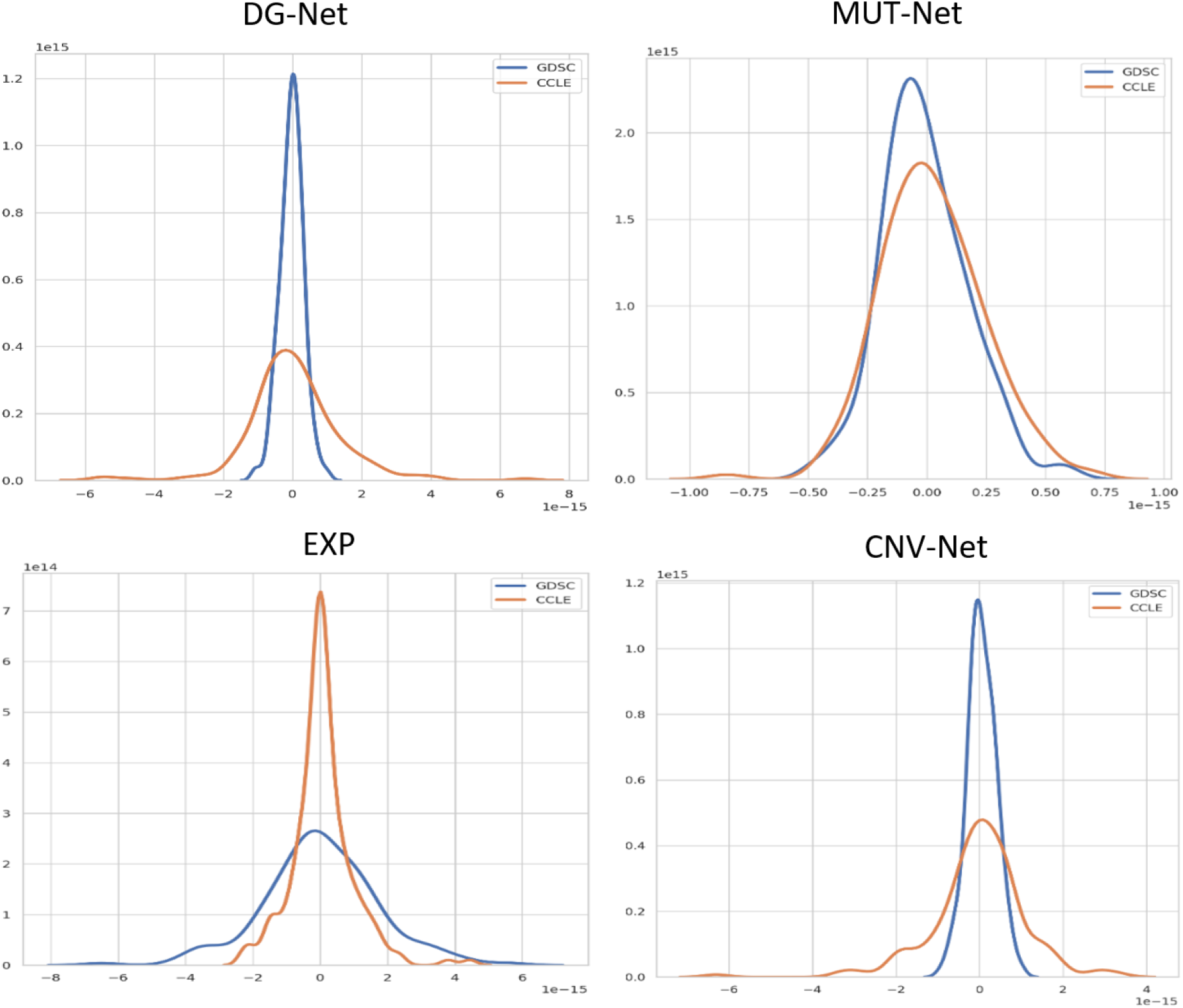
Comparison of enriched pathway feature distribution between the GDSC (blue line) and CCLE (orange line) databases for DG-Net, EXP, Mut-Net and CNV-Net features.

### Explainability of the model

We computed Shapley values to identify important features underlying predictions. We focus on features that have positive correlations with drug response (i.e. the feature contributes to reducing IC50). The top features are distributed across the different data types, including expression, DG-Net, CNV and mutation, supporting our observation that the combination of these data types enables the good performance (figure S1). We highlight a subset of these globally important top features (i.e. features important across all drugs and cell lines in the GDSC dataset): The top feature is an enrichment of cell line expressed genes within the ADP-Ribosylation factor 3 (ARF3) pathway. The main gene of this pathway, ARF3 gene, is up-regulated in breast cancer and promotes breast cancer cell proliferation, representing a novel prognostic marker and therapeutic target for breast cancer (Huang *et al,* 2019) (the GDSC dataset includes 11 breast cancer cell lines).

The second top feature is enrichment of the Histone deacetylase III (HDAC-III) pathway within the drug-associated gene network (DG-Net). HDAC inhibitors seems to be promising anti-cancer drugs (Eckschlager *et al*, 2017). Several genes within this pathway have established role in cancer, such as TP53(Olivier *et al,* 2010; Petitjean *et al,* 2007) and SIRT1, which is up-regulated in cancer cells and may play a critical role in tumor initiation, progression, and drug resistance by blocking senescence and apoptosis, and promoting cell growth and angiogenesis. SIRT1 inhibitors have shown promising anticancer effects in animal models of cancer (Liu *et al,* 2009). The third top feature is CNV of Glypican 1 pathway, further discussed in the leave-one-cell-out specific example below. Finally, the next two features involve enrichment of the Nuclear factor kappa B (NF-κB) canonical pathway within mutation data and enrichment of Fanconi Anemia pathway by DG-Net. Indeed, mutations in the component of the core NF-κB signaling pathway have been implicated with relation to cancer (Courtois & Gilmore, 2006) and patients with Fanconi Anemia have a higher risk of cancer, particularly for acute myeloid leukemia and squamous cell carcinoma followed by ongoing work to study targeting of pathways that are synthetic lethal with loss of Fanconi Anemia pathway with Olaparib or other drugs (Nalepa & Clapp, 2018).

Next, we highlight two local feature importance predictions, one for the LODO (RVX208) and one for the LOCO scenario, demonstrating the drug- and cell line-specific explanability of PathDSP.

#### RVX208

The best performing prediction of LODO is for RVX-208 (apabetalone, RMSE = 0.34 across all cell lines). RVX208 is a Bromodomain and Extra-Terminal domain (BET) inhibitor. BET regulates the transcriptional program and plays a role in influencing cancer pathogenesis and inflammation (Gilan *et al*, 2020). The Activin receptor-like kinase-1 (ALK1) pathway ranked highest out of the features with positive correlation with drug responses (Figure S2). ALK1 is a receptor of TGF-beta type 1 receptor family and can regulate angiogenesis. Angiogenesis plays a critical role in the growth of cancer because solid tumors need a blood supply if they are to grow in size and tumors can actually cause this blood supply to form by stimulating angiogenesis (Nishida *et al*, 2006). Correspondingly, ALK1 is a well-known cancer driver which could act as a tumor suppressor or oncogene, depending on the cancer type, cell type, or ligand involved (Valer *et al*, 2019) and is an emerging target for antiangiogenic therapy of cancer (Cunha & Pietras, 2011). Additional genes in this pathway include mitogen-activated protein kinase 1 and 3 (MAPK1 and MAPK3), where the suppression of BET inhibits vascular inflammation by blocking MAPK activation (Huang *et al*, 2017).

#### Chronic Myelogenous Leukemia cell line

Our model obtained good performance when predicting drug response for the Chronic Myelogenous Leukemia (CML) cell line (SIDM00482) in the LOCO (RMSE = 0.28). Based on the Shapley values, gene expression enrichment of the SYNDECAN 3 pathway and copy number variations enrichment of the GLYPICAN 1 pathway display positive correlation with drug response (Figure S3). Syndecans and glypicans are membrane-bound heparan sulphate proteoglycans (HSPGs), they act as receptors and relate to cell growth and differentiation by interacting with other growth factors (Cat & David, 2001; Cheng *et al*, 2016). Indeed, syndecans and glypicans have reported roles in tumorogenesis in blood cancers (Sebestyén *et al,* 1997). Specifically, Glypican-1 (GPC-1) was found to be overexpressed and a biomarker of certain cancers with demonstrated ability to distinguish between healthy controls and advanced cancer patients with 100% accuracy (Herreros-Villanueva & Bujanda, 2016), while Syndecan-3 has not been implicated in cancer yet (Cheng *et al*, 2016). However, other members of these pathways include epidermal growth factor receptor (EGFR) that have been implicated with regard to activation in CML (Corrado *et al*, 2016), SRC Proto-Oncogene, Non-Receptor Tyrosine Kinase (SRC) that appears in both Gypican-1 and syndecan-3 pathways, whose overexpression have been identified among the known mechanisms of resistance to imatinib in CML (Quintás-Cardama *et al*, 2006) and FYN proto-oncogene, Src family tyrosine kinase that is up-regulated in CML as result of the BCR-ABL1 oncogene (Ban *et al*, 2008).

These two examples demonstrate that identifying cancer pathways association with drug-associated genes can help identify potential mechanisms and possibly new targets within the pathway.

## Discussion

In this study, we presented PathDSP, a method that integrates drug and cell line information to predict drug responses of cancer cell lines. We addressed the high dimensional and heterogeneous nature of the data used in this task by mapping it to curated pathways. By using this approach, we gained improved performance over current state-of-the-art methods, improved generalizability of the models across independent datasets and provided a cancer pathway-level of explainability beyond that of the single gene or single mutation level. While we focused on manually curated cancer pathways and on specific genomics data including gene expression, mutation and copy number variation data, our framework is flexible and can seamlessly integrate additional pathways as well as other data types.

We demonstrated that integrating multiple types of data improve performance of the algorithm. Interestingly, for drug-related data types, relying only on engineered features from the drug-gene-network without chemical structure information obtained the worst performance, but the combination of the two improved performances over only one of them. Similarly, with regard to cell line specific data types, each of the individual cell-line data, i.e. gene expression, mutation and copy number variation, gain similar but reduced performance relative to their combination. Nevertheless, the reduction in performance in this case is smaller, holding a promise that our method could be applied even within clinical settings with only a subset of the data types measured.

The incompleteness of drug target data poses a potential limitation on good performance of PathDSP. In order to address that, we expanded the list of drug targets to include drug-associated genes from curated external databases and additional functionally-related genes by searching close neighbors of drug target genes within a protein-protein interaction network, potentially capturing also cross-talking pathways. As we demonstrate, the differences in response value measurements (IC50) across databases pose a limitation on the generalizability of methods designed to predict the response value, including PathDSP.

However, we managed to improve the generalizability beyond the expected theoretical difference between the datasets (RMSE of 0.95 vs. 1.13 theoretical), demonstrating good generalization capabilities, which would be critical in translating this method into clinical setting.

Last, while pathway activity may be helpful in predicting drug response differences, biological pathway usually consists of several genes responsible for diverse functionality. Thus, more fine-tuned, drug and cell-line specific, experiments are necessary in order to pinpoint which gene(s) may be the ones most associated with a drug’s sensitivity or resistance. Additionally, while the explainable associations are good predictors, they are not necessarily causal, a relationship that would require additional experiments to establish.

## Conclusion

Our developed method of pathway-based deep neural network for drug sensitivity prediction demonstrated improved performance, generalizability and exemplified explainability. Given the flexibility of this approach, we believe it provides a starting point to evaluate the roles of pathways in drug response for cancers and to provide a steppingstone towards cancer precision medicine.

## Methods

### Data

Data sensitivity data, cell-line gene expression, somatic mutation and copy number variation data for 319 cancer cell lines and 153 drugs was downloaded from Genomics of Drug Sensitivity in Cancer (GDSC). Drug sensitivity data of 24 drugs and 478 cancer cell lines from the Cancer Cell Line Encyclopedia (CCLE) were downloaded through the DepMap portal (https://depmap.org/portal/download/, release version: public 20Q1), which also include gene expression, mutation and copy number variation data for those cancer cell lines. Primary target data was downloaded from GDSC, and PID pathways were downloaded from MSigDB (Liberzon *et al,* 2011). Protein-protein network was downloaded from STRING database (Szklarczyk *et al,* 2019), including protein interactions, co-expression and text-mined interactions.

### Feature engineering and normalization

#### Gene expression data

Gene expression data were measured by transcripts per million (TPM) and log-transformed. We imputed with mean for the rest of missing values. Enrichment score (ES) of each PID pathway in each cell line was calculated using the single-sample Gene Set Enrichment (ssGSEA) algorithm (Barbie *et al,* 2009) through GSEApy (https://gseapy.readthedocs.io/en/master/gseapy_example.html). We ran permutation test for 1000 times and normalized ES scores by the size of gene set to obtain normalized ES (i.e., NES). We used the resulting pathway enrichment matrix with size of 319 cancer cell lines by 196 pathways as the gene expression feature (EXP).

#### Somatic mutation data

Mutation data are in long form, in which each row consists of a cancer cell line and its mutated gene name. We collected all mutations for each cell line to perform network-based pathway enrichment analysis by the NetPEA algorithm (Liu & Ruan, 2013), which calculates an enrichment score by measuring the closeness of pathway genes to a given gene set within a protein-protein interaction (PPI) network. We implemented the algorithm with some modification. The method is summarized as follows: First, both mutation gene and pathway genes were mapped to STRING PPI network (Szklarczyk *et al*, 2019). For each cell, its mutation gene set is then used as the restart nodes to diffuse information through edges to their neighbors within the PPI by the Random Walk with Restart approach. For each pathway, a similarity score between the mutation gene set to the pathway genes is calculated by averaging all values on the pathway genes, where the node value represents the probability of being revisited (i.e., the closeness to the restart nodes). We modified the similarity score by multiplying each gene score with its gene expression value within the cell line, forming a cellspecific gene co-expression-mutation network. Then, a permutation test was performed 1,000 times by randomly selecting the same number of genes, resulting in 1,000 association scores as the background for the pathway. Pathway significance was normalized using z-score, resulting in pathway enrichment matrix with size of 319 cancer cell lines by 196 pathways for the mutation feature (MUT-Net).

#### Copy number variation data

Copy number variation data were represented by the GISTIC (Genomic Identification of Significant Targets in Cancer) score comprising of −2 (deletion), −1 (loss), 0 (diploid), 1 (gain), and 2 (amplification), genes with GISTIC score of 0 were excluded. For each cell line, we collected a set of genes with copy number variations to calculate pathway enrichment score using the same procedure used for mutation data, resulting in the pathway enrichment matrix with size of 319 cancer cell lines by 196 pathways for the copy number variation feature (CNV-Net).

#### Drug gene network data

The primary target genes were provided in the GDSC data, we further expanded the target genes by two approaches. First, we obtained off-target genes from the DGIdb version 3 database (Cotto *et al*, 2018), a webserver collected curated drug-gene interaction data from literature. Second, we used the expanded gene list to find its neighbors within the STRING PPI network (Szklarczyk *et al*, 2019), which increases the number of related gene per drug from 4.71 to 876.78 on average. For each drug, we performed pathway enrichment analysis against 196 PID pathways (Schaefer *et al,* 2009) using its expanded target gene list. We used the original NetPEA approach described by Liu et al., (Liu & Ruan, 2013). i.e., we did not aggregate gene expression value to the node like we did for mutation and copy number variation data. As a result, we obtained the pathway enrichment scores in drug targets of size 144 drugs by 196 pathways for the drug gene network feature (DG-Net), excluding nine drug without target information or the target was missing from the PPI network.

#### Chemical structure data

We retrieved canonical SMILE strings by searching the PubChem database (Kim *et al,* 2016) with the open-source Python API, PubChemPy (https://pubchempy.readthedocs.io/en/latest/). We then converted SMILE strings into Morgan fingerprint with the open-source cheminformatics toolkit RDKit (http://www.rdkit.org), generating the matrix of 153 drugs by 256 molecular bits for the chemical structure feature (CHEM).

### Model fitting

We created a fully connected neural network (FNN), following an architecture suggested by Li et al., (Li *et al,* 2019) using Pytorch (Paszke *et al,* 2019), parameters used is listed in the Table S2. We performed 10-fold cross validation for the experiments in this study with early stopping applied to avoid overfitting, repeated five times for robustness verification. Before feeding into the FNN model, we normalized our features using z-score. For the experiment of leave-one-drug-out, we took out one drug from training each time (with all its cell-lines), and for the experiment of leave-one-cell-line-out, we took out one cellline from training (with all its drugs). We used RMSE to estimate generalization error of the model. For compared machine learning algorithms besides FNN, we used scikit-learn API (Buitinck *et al,* 2013) for the other models, including the XGBoost algorithm with hyperparameter tuning and early stopping.

### Generalization assessment against CCLE

We tested the generalizability of PathDSP by training on GDSC and applying the trained model to the CCLE dataset. We then calculated RMSE for all samples and additionally for six drugs and 35 cancer cell line pairs shared between the CCLE and GDSC datasets.

### Feature importance

Feature importance was measured as the amount of contribution each feature makes to the prediction value. We used the Shapley value (Shapley, 1953) to estimate feature importance, which is measured by comparing the prediction value obtained with the feature and without it (Lundberg & Lee, 2017). The python library SHapley Additive exPlanations (SHAP) (Lundberg *et al,* 2018) was used to obtain global feature importance and visualization of feature importance at the local level. If a feature X has a positive shapely value, it indicates that feature X contributes to a higher predicted value, and vice-versa. Thus, in this study, it is interpreted as the feature X increases/decreases drug responses to drug Y (i.e. increase in −log(IC50), which means decrease in IC50).

## Code availability

PathDSP code is available at: https://github.com/TangYiChing/PathDSP

## Data availability

All relevant data used in this study is publically available and detailed in the data section of the manuscript.

## Author contributions statement

Y.T. conceived and conducted the experiment(s), Y.T. and A.G. analyzed the results. All authors reviewed the manuscript.

